# Paternal germ line aging: DNA methylation age prediction from human sperm

**DOI:** 10.1101/220764

**Authors:** Timothy G Jenkins, Kenneth I Aston, Andrew Smith, Douglas T Carrell

**Affiliations:** Andrology and IVF Laboratories, Department of Surgery; Department of Obstetrics and Gynecology; Department of Genetics, University of Utah School of Medicine, Salt Lake City, Utah, USA; University of Southern California

**Keywords:** Sperm Epigenetics, aging, DNA methylation, aging calculator

## Abstract

**Background:** The relationship between aging and epigenetic profiles has been highlighted in many recent studies. Models using somatic cell methylomes to predict age have been successfully constructed. However, gamete aging is quite distinct and as such age prediction using sperm methylomes is ineffective with current techniques.

**Results:** We have produced a model that utilizes human sperm DNA methylation signatures to predict chronological age by utilizing methylation array data from a total of 329 samples. The dataset used for model construction includes infertile patients, sperm donors, and individuals from the general population. Our model is capable of accurately predicting age with an R2 of 0.928 in our test data set. We additionally investigated the repeatability of prediction by processing the same sample on 6 different arrays and found very robust age prediction with an average standard deviation of only 0.877 years. Additionally, we found that smokers have approximately 5% increased age profiles compared to ‘never smokers.’

**Conclusions:** The predictive model described herein was built to offer researchers the ability to assess “germ line age” by accessing sperm DNA methylation signatures at genomic regions affected by age. Our data suggest that this model can predict an individual’s chronological age with a high degree of accuracy regardless of fertility status and with a high degree of repeatability. Additionally, our data appear to show age acceleration patterns as a result of smoking suggesting that the aging process in sperm may be impacted by environmental factors, though this effect appears to be quite subtle.

## INTRODUCTION

In the very recent past a great deal of work has been performed in an effort to understand the nature of aging, the mechanisms that drive the process, and the biomarkers that may be predictive of, or affected by, age. In this effort, a seminal manuscript was published in 2013 which described the ability to use DNA methylation signatures in somatic tissues to predict an individual’s chronological age [1]. In this work, Dr. Horvath demonstrated that the epigenetic mechanisms that reflect the aging process are tightly conserved between individual tissues and across multiple species. Remarkably, these patterns are sufficiently consistent to enable accurate age prediction with Horvath’s age calculator despite the significant contrast in epigenetic profiles between various somatic tissues.

Despite the general applicability of this model across diverse tissues, one tissue in particular did not display similar predictive power as was seen with most. In fact, DNA methylation signatures from testicular tissue and sperm specifically did not appear to be predictive of age at all with the previously described calculator [1]. In agreement with this observation is data from our lab which suggests that the nature of age associated alterations to sperm DNA methylation signatures are opposite of what is typically seen in somatic cells [1-4]. Specifically, although aging results in a global decrease in methylation and increased regional methylation in most cell types, we demonstrated that sperm exhibits the opposite trend. In many ways such a finding is not surprising as this is not the first case where the male germ line defied conventional age-associated cellular alterations. The most well described example of this is that of age impacts on telomere length. A hallmark of aging in somatic cells is a marked shortening of telomeres, but in sperm telomere lengthening is commonly seen with aging [5]. Clearly, sperm cells are extraordinarily unique and thus it seems likely that a unique approach is required to understand both the nature of the aging process and the potential predictive power of age associated alterations to the sperm epigenome.

In our previous publications we have described the general impact of aging on the sperm methylome. In these studies, we have shown that sperm have a very distinct pattern of age-associated alteration [2, 3]. We identified 148 genomic regions (~1kb in size) that displayed differential methylation with age. Of these, only 8 displayed an increase in methylation, and the remaining 140 regions experienced a marked loss of methylation with age. Intriguingly, these regions of differential methylation are enriched at genes known to be associated with bipolar disorder and schizophrenia, both diseases known to have increased incidence in the offspring of older fathers. Indeed the epigenetic patterns of aging in sperm, while distinct from the epigenetic patterns of aging in somatic tissues, are striking and extremely consistent and thus provide an excellent opportunity for predictive model construction.

The pursuit of generating a model to predict an individual’s age using the sperm methylome is not only an interesting question from the perspective of basic cell biology but the patterns of sperm aging, and the unique nature of the sperm make the utilization of this cell type ideal for such a predictive model. Using pure cell populations is ideal for any epigenetic analysis, and while the previously constructed models are effective at predicting age even with tissues that are difficult to purify (which is a testament to quality of model and to the strength of the aging signal), the ideal scenario would be to use a pure cell population. Human sperm offer just such an opportunity. Many protocols are applied to somatic cell removal in sperm epigenetic studies and they have proven quite effective at isolating only germ cells, thanks in large part to the highly unique and compact nature of the sperm head. Further, the magnitude of the aging signal is quite strong in the sperm (thought to be in part due to the highly proliferative nature of the sperm cells themselves) and as a result, the patterns of aging offer an excellent opportunity for powerful prediction. In this study, we set out to capitalize on these advantages to build a model that can predict an individual’s age using methylation signatures in the paternal germ line. The experiments outlined herein describe the utility of the germ line age calculation and also provide evidence to suggest that the rate of aging can be affected by environmental exposures or lifestyles (smoking, obesity, etc.).

## RESULTS

### Model construction and training

In the current study we assessed sperm DNA methylation array data (Illumina 450K array) from 3 distinct previously performed studies [2, 6, 7]. From these data sets, we were able to utilize a total of 329 samples that were used to generate the predictive model outlined herein. Individuals with many different fertility phenotypes provided the samples used in this study. Specifically, our training data set includes samples from sperm donors [2], known fertile individuals, infertility patients (including those seeking intrauterine insemination or even in vitro fertilization treatment at our facility), and individuals from the general population [6, 7]. Further, our data set includes those that have very different lifestyles and environmental exposures (as an example, both heavy smokers and never smokers are represented in our data set).

We utilized the glmnet package in R to facilitate training and development of our linear regression age prediction model [8]. For training of our model, we limited the training dataset to only the 148 regions previously identified to be strongly associated with the aging process to ensure clear interpretability of the model [2]. Beta-values were used in all experiments. These values represent fraction methylation as the standard output from the Illumina methylation array, which are scored between 0 and 1 with 0 representing complete demethlyation and 1 representing complete methylation. We designed multiple iterations of test scenarios by adjusting the features used to achieve the most predictive power. First, we trained on all of the beta-values for each CpG located in our regions of interest (“CpG level” features). Second, we generated a mean of beta-values for each region, which included the CpGs within each region respectively. Ultimately, this approach yielded mean beta-values for each region (“regional level” features), and the model was trained only on these averages.

In each of the above-described scenarios, we employed a 10-fold cross validation strategy. This was performed 10 times on unique subgroups of the entire data set (Figure 1A-F). The results from these ten validations were compared between the CpG level training and the regional level training. To compare the accuracy and predictive power of these models we performed linear regression for each (actual age vs. predicted age) and generated R^2^ values. These R^2^ values were compared via simple two-tailed t-test to determine if any significant differences exist between the two approaches to model construction (CpG level construction vs. regional level construction). These tests revealed that there was a modestly significant decrease in predictive power in the regional model when considering only the training data sets (p=0.0428). However, there was no significant difference seen between the test sets (p=0.3439). In fact, while not significant, the CpG level model appeared to be more prone to extremes in significantly lower predicative power in individual test sets when compared to the regional level models (Figure 1G). In an effort to make the model as simple as possible and in light of these findings, we committed to use the regional level model moving forward. Further, the alterations that occur at single CpGs are less likely to be biologically meaningful than those that occur over a region of the genome, thus using the regional level feature set improves the biological interpretability.

**Figure 1:**
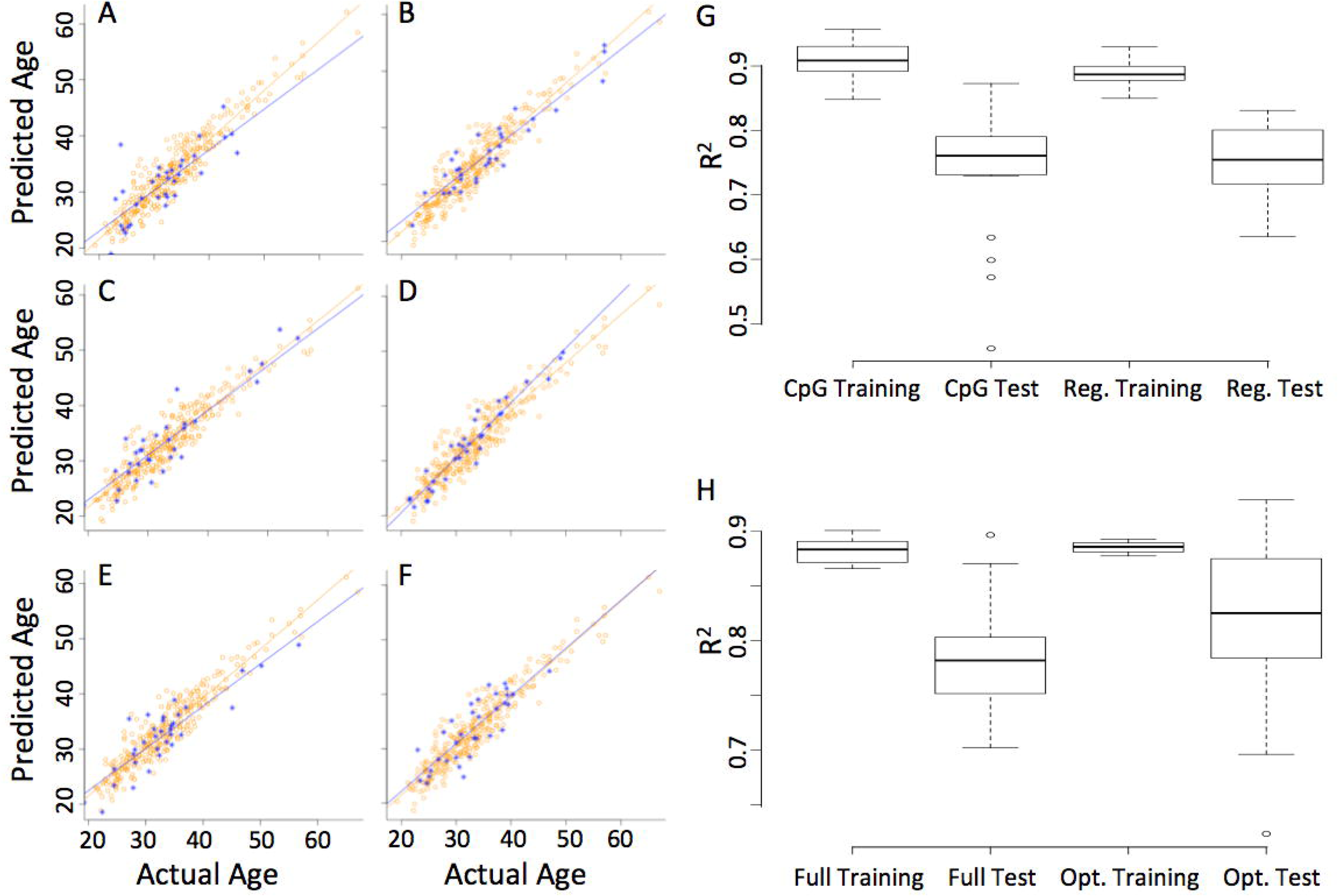
(A-F) Scatterplots depicting the relationship between predicted and chronological age in 6 represented models from our cross validation testing. (G) Box and whisker plots of the R2 values from each cross validation (10) for both training and test datasets between the CpGs level data and the regionalized data. (H) Box and whisker plots of the R2 values from each cross validation (10) for both the full regional data set (147 regions) and the optimized regional data set (51 regions) with both training and test data displayed.

We additionally assessed the weighting of the features (regions) used in the models constructed during cross validation. We found a great deal of variation in the features selected across the regions screened, though a subset of the regions were heavily weighted and used in 80% or more of the models built during cross validation (a total of 51 features/regions met this criterion). In an effort to identify the simplest model we compared cross validation (10-fold strategy) in only these 51 regions (“optimized regions”) to all of the regions previously screened. We found that both the training and test groups were not statistically different between the optimized regional list and the full regional list (Figure 1H). We therefore selected the best performing model that was trained only on 51 regions of the genome (Table 1). With this model we are able to generate a prediction of all 329 samples in our data set with an r^2^ of approximately 0.89 and an average accuracy in prediction of 93.7%.

**Table 1:**

| **Name** | **CHR** | **Start** | **Stop** |
| --- | --- | --- | --- |
| ADAMTS8 | chr11 | 130299298 | 130299948 |
| ARC | chr8 | 143694010 | 143694548 |
| ARGHGEF10 | chr8 | 1877888 | 1878324 |
| BCL11A | chr2 | 60680616 | 60680762 |
| C1ORF122 | chr1 | 38272200 | 38273057 |
| C7ORF50 | chr7 | 1083209 | 1084163 |
| CCDC144NL | chr17 | 20798895 | 20799770 |
| CLIC1 | chr6 | 31698492 | 31699299 |
| DMPK | chr19 | 46282571 | 46283081 |
| FAM86C1 | chr11 | 71498202 | 71499118 |
| FAM86JP | chr3 | 125634060 | 125634453 |
| FOXK1 | chr7 | 4722778 | 4723928 |
| FSCN | chr7 | 5635134 | 5635954 |
| GAPDH | chr12 | 6641602 | 6642355 |
| GET4 | chr7 | 914964 | 915832 |
| GNB2 | chr7 | 100274361 | 100275305 |
| GPANK1 | chr6 | 31630819 | 31632542 |
| GPR45 | chr2 | 105857809 | 105859084 |
| KCNQ1 | chr11 | 2554562 | 2555577 |
| LDLRAD4 | chr18 | 13611370 | 13611825 |
| LMO3 | chr12 | 16760040 | 16761003 |
| LOC100133461 | chr4 | 3680721 | 3681760 |
| MIR22HG | chr17 | 1617363 | 1618296 |
| MTMR8 | chrX | 63614857 | 63615496 |
| N10 | chr1 | 28423399 | 28424202 |
| N12 | chr5 | 3593413 | 3594276 |
| N22 | chr19 | 4579481 | 4580471 |
| N23 | chr14 | 106004434 | 106004608 |
| N24 | chr6 | 170449417 | 170450804 |
| N27 | chr6 | 30432200 | 30433944 |
| N30 | chr15 | 27959473 | 27960032 |
| N8 | chr11 | 69260136 | 69261045 |
| N9 | chr7 | 35300077 | 35301070 |
| NCOR2 | chr12 | 124990897 | 124991140 |
| NONE | chr10 | 17347047 | 17347392 |
| NSG1 | chr4 | 4386726 | 4387698 |
| PAX2 | chr10 | 102509693 | 102510569 |
| PITX1 | chr5 | 134365728 | 134366535 |
| PRSS22 | chr16 | 2908157 | 2908935 |
| PTPRN2.3 | chr7 | 157523356 | 157524159 |
| PTPRN2.4 | chr7 | 158109339 | 158110153 |
| PURA | chr5 | 139492535 | 139493491 |
| PYY2 | chr17 | 26553567 | 26554908 |
| SECTM1 | chr17 | 80278592 | 80280331 |
| SEMA6B | chr19 | 4555999 | 4556983 |
| SEZ6 | chr17 | 27330794 | 27332647 |
| SLC22A18AS | chr11 | 2909690 | 2909716 |
| SOHLH1 | chr9 | 138590204 | 138590996 |
| THBS3 | chr1 | 155176868 | 155177784 |
| TNXB | chr6 | 32064146 | 32065891 |

### Technical validation / replicate performance

Because variability can be a concern in array experiments, we tested our model in a completely independent cohort of samples that were not used in any of our cross validation / model training experiments. We utilized 10 sperm samples, each with 6 replicates (a total of 60 samples) that were each run on the 450K array platform from a previously published study [9]. Further, the samples from this study were exposed to varying extremes in temperature to test the stability of the sperm DNA methylation signatures. Thus these samples do not represent strict technical replicates (because of slight variations in treatment) but do provide an even more robust test of the algorithms predictive power on sperm DNA methylation signatures in multiple samples from the same individual. The model was applied to these samples and performed well with the average standard deviation in age prediction being only 0.877 years. We additionally performed linear regression analysis on the average predicted age vs. actual age in each of the 10 individuals in the dataset and found a significant association between these two (R^2^ of 0.732; p=0.0016; Figure 2).

**Figure 2:**
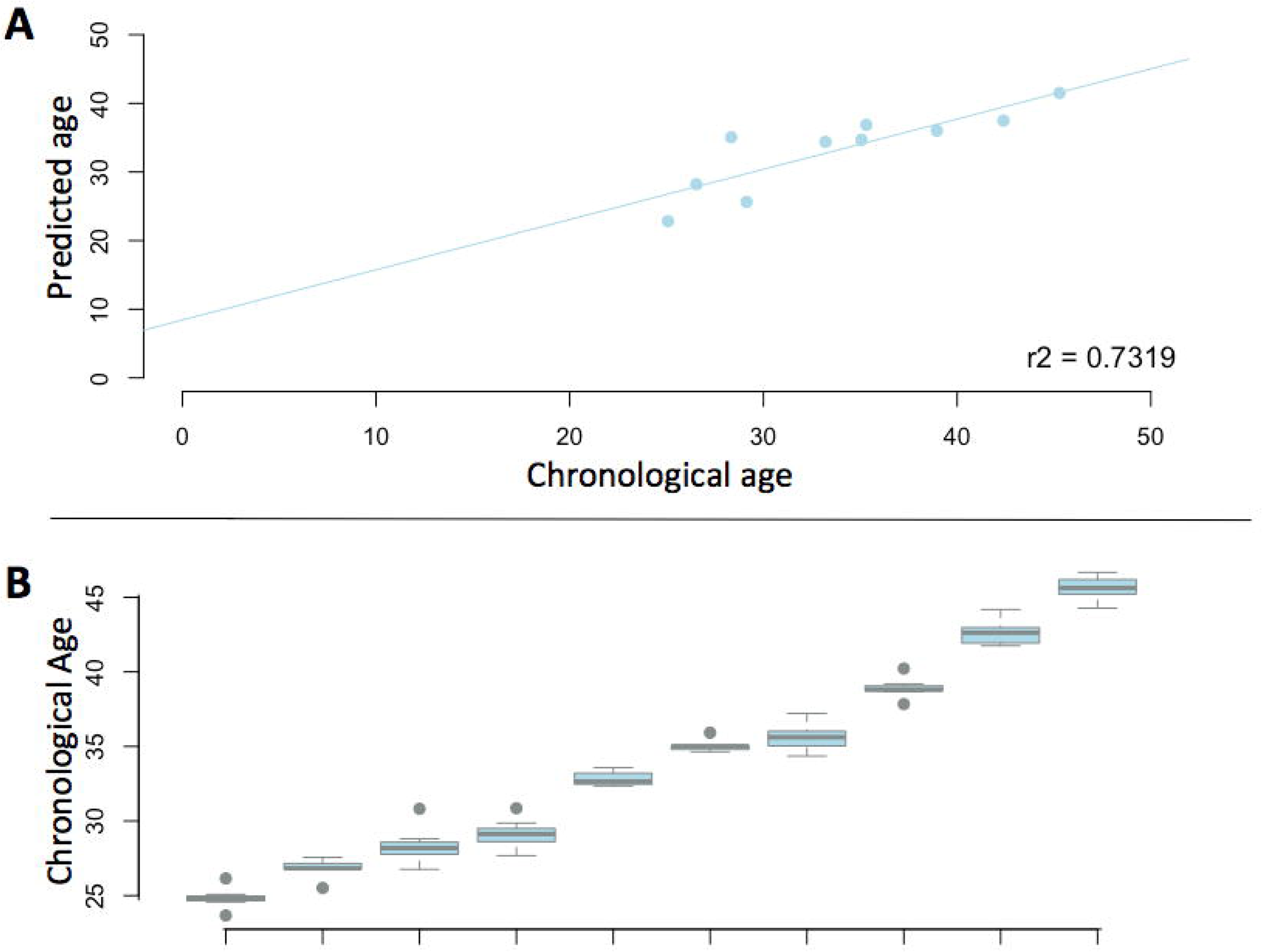
(A) scatterplot depicting the age prediction in a completely independent cohort of samples. (B) Boxplots demonstrating the variation in age prediction from ten individuals with six biological replicates that were run in a completely independent cohort.

### The impact of smoking on age prediction

To test the potential diagnostic/clinical utility of our model we have more closely assessed the data in our original cross validation dataset. Specifically we have analyzed our smoking dataset [7], which includes sperm methylation data from 78 smokers and 78 individuals who responded as “never smokers.” Similar aged men are represented in each group. We additionally isolated a portion of the smoking group who were had smoked cigarettes for >10 years. We found an approximately 1.5% increased in predicted age compared to chronological age in all smokers and 2.5% increase in long term smokers. However this difference failed to reach statistical significance. Interestingly, this same pattern was observed (though significantly higher in magnitude) when screening only individuals who were less than 35 years old at the time of collection (Figure 3). In these samples we saw a 3% increase in predicted age compared to chronological age in the smoker group and a nearly 6% increase in predicted age in the long-term smokers (p=0.0196).

**Figure 3:**
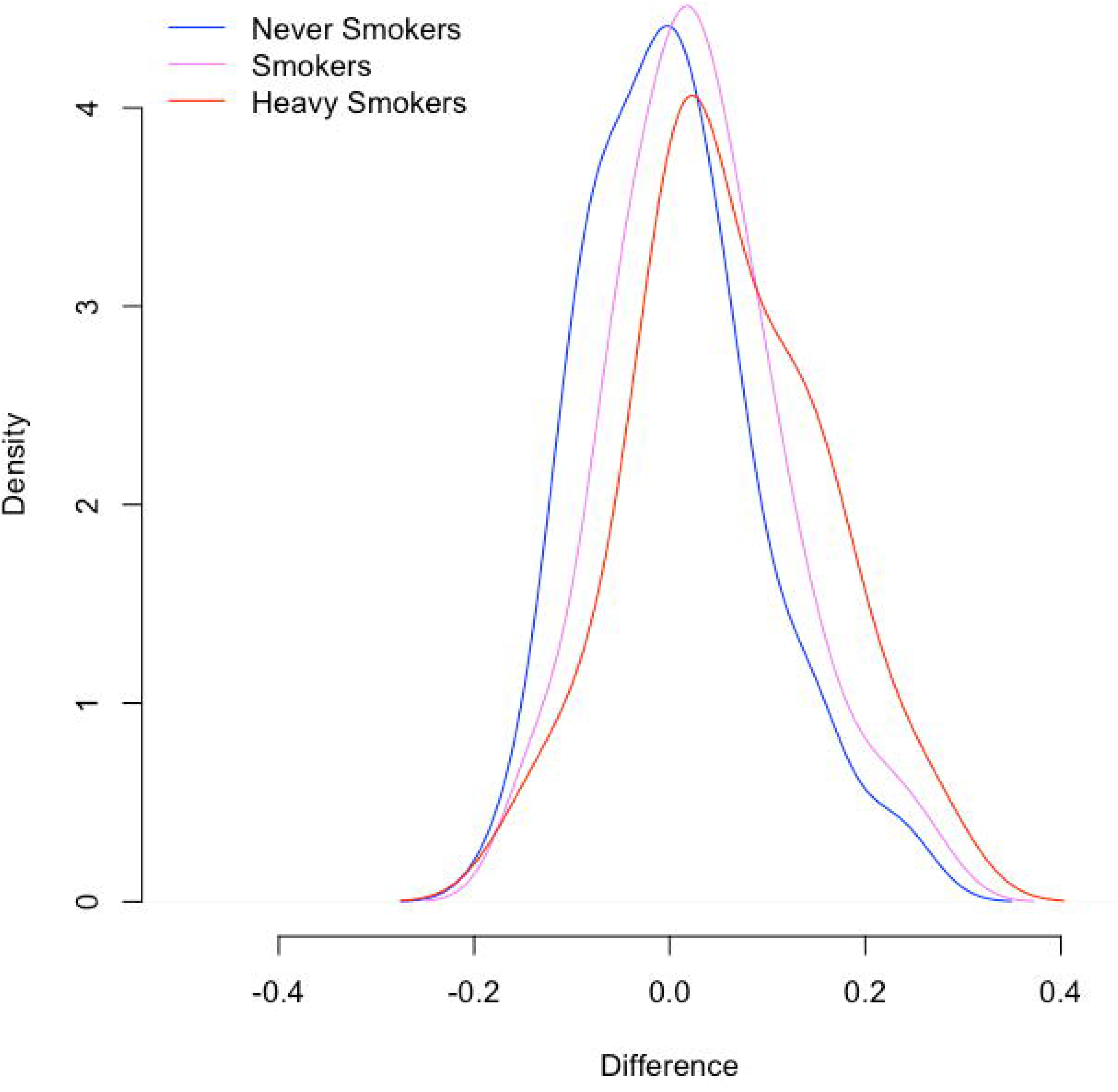
Density plot shows the accuracy of age prediction in never smokers, smokers, and heavy smokers among individuals below 35 years of age. Similar patterns exist in the entire cohort but are the most profound in this age group.

## DISCUSSION

We have developed a sperm age calculator that has the capacity to identify an individual’s chronological age based only on their sperm DNA methylation signatures. Previous studies have defined aging patterns in somatic cells and one in particular [1] very successfully generated an aging calculator using methylation signatures of somatic cells. However, these findings do not hold true in sperm and further, the DNA methylation age calculator that described in 2013 fails to work effectively with paternal germ line epigenetic signatures. Herein, we have described the development of a linear model that has the ability to accurately predict ages with these signatures. Specifically our model is based on average methylation signatures at 51 genomic loci known to be altered as men age [2].

In the process of model construction, we evaluated multiple potential methods by which we could train our model. One important consideration was the nature of the population with which the model was trained. While there is a balance in selecting a population (broad applicability vs. targeted population) we decided to utilize a population with diverse fertility phenotypes and exposures to ensure that it could perform well with many different phenotypes. As such we included smokers and non-smokers, individuals of known fertility, those currently being treated for infertility, and men from our general population.

We also attempted to obtain a simple model. While training on all data from the entire array may have provided additional power in prediction, it also would likely make the model very difficult to interpret. Instead, we focused only on the regions that we knew were independently altered by age (based on previous data) and refined the model by only assessing these regions. We found that even in our model using regional level features there was some amount of simplification that could be performed. Indeed, we were able to scale our list of features from 148 regions down to 51 regions with the same predictive power. This effort resulted in a quite robust model with strong predictive power (an average of ~94% accuracy in predicting each indiviudals age and an R2 of ~0.89). Had our final model required a great deal of improvement, there would be a need to revisit our targeted approach by increasing our feature set to include more, or all, of the array. However, since we have such a robust and interpretable model as it stands, pursuing a different course was not warranted.

Our data indicate that the model constructed herein is also technically robust. We were able to assess previous data from our lab in which 10 individuals had six technical replicates on 450k methylation arrays [9]. This replicate data enabled to assess the power of the model in two distinct ways. First, we were able to assess the predictive power of the model on a completely independent cohort (each of these samples were assessed at a different time and in different batches of arrays than those upon which the model was trained). Second, we were able to show that the model is able to generate consistent predictions for individuals between technical replicates. Of additional interest is the fact that the samples used in these technical replicates originated from a study that tested the impact of extreme and prolonged temperature exposures on sperm DNA methylation patterns. Thus a portion of the replicates screened were exposed to various magnitudes of less than ideal conditions. However, it is important to note that we did observe a drop in R^2^ in our independent cohort. This is not an entirely unexpected finding due to the variation that can occur between different batches of arrays. In brief, we found that the array batch effects are sufficiently strong to slightly decrease the predictive power of the age calculator for batches performed outside of our original training set. In contrast, the model is sufficiently strong to overcome such variation with strong predictive power, though that power is slightly reduced compared to what is seen in our training/test data set. The ability to maintain predictive power, even when assessing other batches of data is important in a model that will have broad applicability. In the future as more data become available, the model can be updated with increased sample size and with additional batches of experiments, which will lead to even more robust predictive power.

Our data also suggest that there may be some utility for such a model in a clinical setting. Specifically, we were able to identify an age-affect of smoking in our cohort of patients.

We found that individuals who smoke appeared to have acceleration in the pattern of aging and thus the individual’s germ line age was in some cases significantly higher than their chronological age. This represents one example of many different analyses that could be performed in a clinical setting. With future studies we may find that different levels/types of infertility, obesity, or other environmental exposures may cause acceleration in the aging pattern seen in sperm. One of the biggest questions that remains if such associations exist is the potential impact of this age acceleration. Such a pattern could potentially result in increased risk to offspring health, as epidemiological data clearly shows increased incidence of neuropsychiatric disease in the offspring of older fathers [10-15]. This increase in risk may not mean that the altered methylation pattern itself causes these offspring abnormalities, but instead the methylation signatures of age are simply a good indicator of the overall state or age of the sperm. Likely of more immediate interest to clinicians is the fact that advanced paternal age is associated with a loss of fecundity and fertility. Specifically, it has been shown that men older than 45 years take ~5 times longer to achieve a pregnancy as men less than 25 years (when controlling for female age) [16]. A similar decrease in fecundity was identified in a large population study in 2000 which showed that (after adjusting for maternal age) men > 35 years of age had a 50% lower chance of achieving a pregnancy within 12 months of attempting conception than younger men [17]. Other studies have also shown decreased fertilizing potential in both IUI and IVF [18, 19]. While the magnitude of this effect remains controversial [20, 21], it is clear that advanced paternal age does play an important role in a couple’s fertility status and can clearly result in, at a minimum, a significantly increased time to pregnancy. For many couples, such potential barriers to achieving a pregnancy are essential to understand and discuss with their care providers. While none of these associations have been proven in this specific work, the potential clinical utility of the calculator is clear and warrants further investigation both in predicting an individual’s health/fertility as well as in the prediction of abnormalities in the offspring.

It is important to note that while the findings of alterations associated with age in the sperm epigenome are intriguing, the direct impact of these alterations is still in question. In fact, the actual impact of any sperm epigenetic alteration on the embryo or the offspring is difficult to predict due to massive reprograming events that take place in the early embryo and in the primordial germ cells. However, data do suggest that methylation marks in many sub-telomeric regions escape reprograming events and can be potentially be passed on to the offspring [22-26]. Intriguingly, our original sperm aging study showed that the majority of age-affected regions were located in these sub-telomeic regions as well [2]. Such a transmission of age-affects would be remarkable, but may offer a real potential explanation for at least a portion of the downstream impact of paternal age on offspring disease incidence and phenotype.

The data described herein are quite promising, though some limitations are clear. Foremost among them is our knowledge of downstream impacts as described above. This will require a great degree of effort to determine the nature of these effects and if risks to fertility or the offspring can be modified in any way by various treatments. Further, while the current model is very effective at predicting an individuals age and is quite robust technically, the alterations we are observing to predict age are subtle and thus small inefficiencies can result in an inability to detect meaningful changes. Despite this, because of the approach we have taken in designing a model based only on limited numbers of regions there is a potential to modify this model for use with different platforms that may offer increased resolution and consistency, for example targeted sequencing [27]. With such an approach, we may be able to improve an already robust predictive model by multiplex sequencing with extreme depth at only the 51 sites of interest. This could provide an even more economical and reliable predictive model. Taken together, the data that we have shown here are intriguing and warrant a great deal of further investigation and also have the potential to be improved with future iterations.

## METHODS

### Samples, study design, data availability

In the current study we assessed sperm DNA methylation array data from 3 distinct previously performed studies [2, 6, 7]. All of the studies were performed in our laboratory. We included only the samples for which ages were available. From these data sets, we were able to acquire a total of 329 samples that were used to generate the predictive model outlined herein. Each sample was run on the Illumina 450K methylation array. In each case, we used SWAN normalization to generate beta-values (values between 0 and 1 that represent the fraction of a given CpG that is methylated) that were used in our study. During early processing of the sperm samples, great care was taken to ensure that no somatic cell contamination was present that could potentially influence the results of our studies. To confirm the absence of somatic cell contamination we assessed the methylation signatures at a number of sites throughout the genome, each of which are highly differentially methylated between sperm and somatic tissues. In Figure 4, we show the differential methylation at one representative genomic locus, DLK1, to illustrate the absence of contaminating signals in the samples used in our study. While variability exists between the methylation in these samples there exists very little, if any somatic DNA methylation signals.

**Figure 4:**
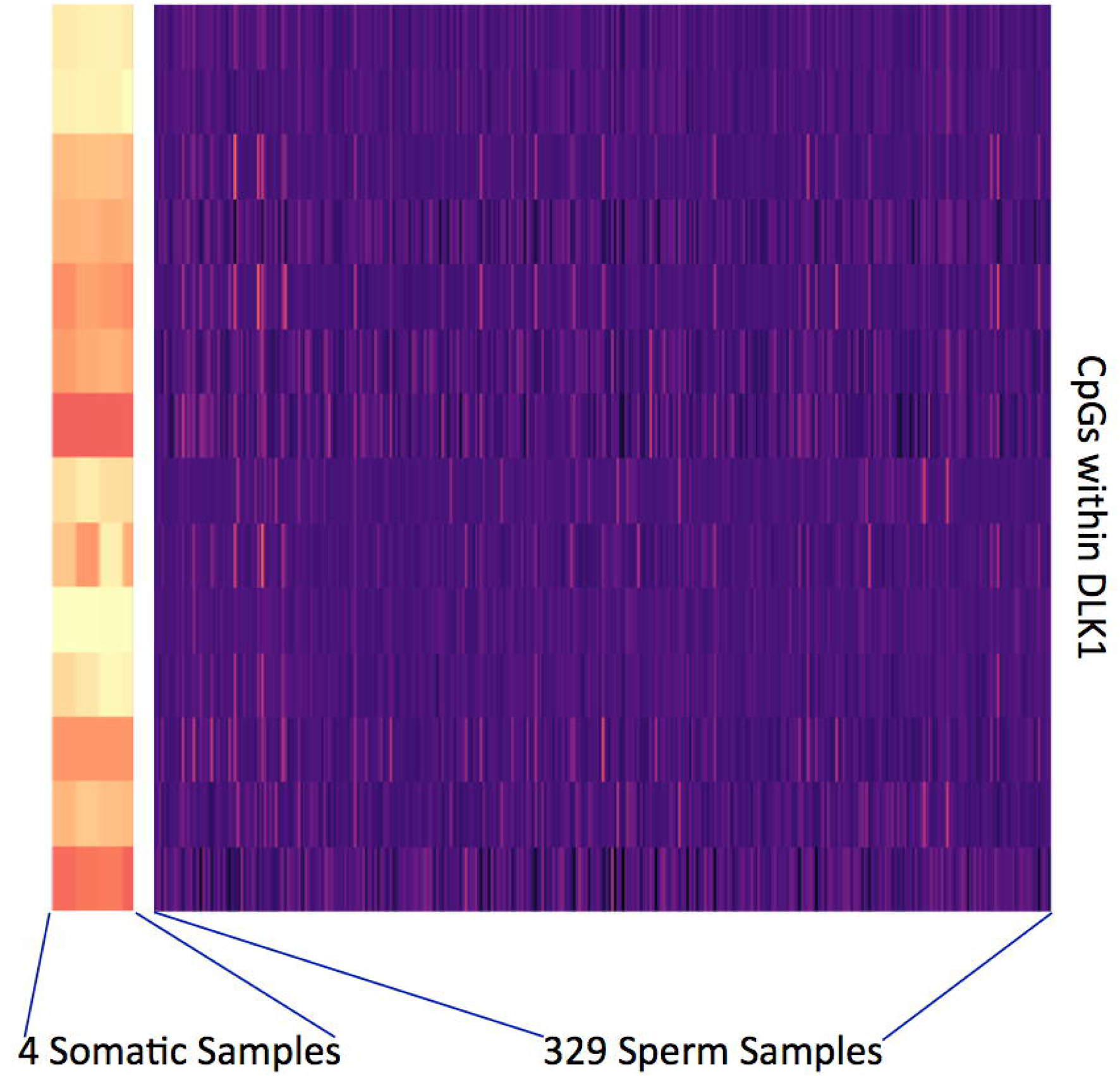
Heatmap of the DLK1 locus, which is highly differentially methylated between sperm and somatic cells is used to confirm the absence of contaminating signals in our data set. 4 blood samples are listed at the far left of the heatmap and the remainder of the samples used in our study follow.

### Samples used

Individuals with many different fertility phenotypes provided the samples used in this study. Our training data set includes samples from sperm donors, known fertile individuals, infertility patients (including those seeking intrauterine insemination or even in vitro fertilization treatment at our facility), and individuals from the general population. Further, our data set includes those that have very different lifestyles and environmental exposures (as an example, both heavy smokers and never smokers are represented in our data set).

### Model Training

We utilized the glmnet package in R to facilitate training and development of our linear regression age prediction model [8]. For training of our model, we limited the training dataset to only 148 regions that we have previously identified to be strongly associated with the aging process to ensure the broad interpretability to the results of the model [2]. We trained two models to identify the best possible outcomes. First, we trained on all of the beta-values for each CpG located in our regions of interest (“CpG level” training). Second, we generated a mean of beta-values for each region that included the CpGs within each region respectively yielding mean beta-values for each region (“regional level” training), and the model was trained only on these averages.

In both of the above-described scenarios, we employed a 10-fold cross validation strategy to repeatedly test trainings on 90% of our samples and hold out 10% for a test set. This was performed 10 times on unique subgroups of the entire data set. The results from these ten validations were compared between the CpG level training and the regional level training. To compare the accuracy and predictive power of these models we performed linear regression for each (actual age vs. predicted age) and generated r^2^ values. These r^2^ values were compared via simple two-tailed t-test to determine if any significant difference exists between the two approaches to model construction (CpG level construction vs. regional level construction).

### Technical validation / replicate performance

We tested our model in a completely independent cohort of samples [9]. We used 10 sperm samples each with six technical replicates that were each run on the 450K array (not those used in our cross validation / model training) to determine the precision and consistency of prediction. Linear regression analysis of predicted vs. actual age was performed using R.

### The impact of smoking on age prediction

We tested 78 never smokers and 78 smokers using our age prediction model. Similar aged men are represented in each group. We additionally isolated a portion of the smoking group who have smoked cigarettes for >10 years. In this analysis we compared accuracy of the age prediction of each group to determine if there is a significant increase in the age prediction compared to chronological age in individuals who smoke. We identified the percent difference between chronological age and predicted age and compared this value between smokers and non-smokers via two-tailed t-test to identify the presence of age acceleration.

## ACKNOWLEDGMENTS

We recognize the efforts of Chris Conley from the Huntsman Cancer Institute for his technical assistance.

## AVAILABILITY

The model generated in this manuscript as well as all instructions on use are publically available on a publically available repository:

Jenkins, TG, Sperm Aging Calculator, (2017), GitHub repository, https://github.com/timgjenkins/Jenkins-et-al-2017

